# Dynamics of spike time encoding in the olfactory bulb

**DOI:** 10.1101/2022.06.16.496396

**Authors:** Jesse C. Werth, Matthew Einhorn, Thomas A. Cleland

## Abstract

In the mammalian olfactory bulb (OB), gamma oscillations in the local field potential are generated endogenously during odor sampling. Such oscillations arise from dynamical systems that generate organized periodic behavior in neural circuits, and correspond to spike timing constraints at fine timescales. While the cellular and network mechanisms of gamma oscillogenesis in the OB are reasonably well established, it remains unclear how these fine-timescale dynamics serve to represent odors. Are patterns of spike synchrony on the gamma timescale replicable and odor-specific? Does the transformation to a spike-timing metric embed additional computations? To address these questions, we used OB slices to examine the spike timing dynamics evoked by “fictive odorants” generated via spatiotemporally patterned optogenetic stimulation of olfactory sensory neuron axonal arbors. We found that a small proportion of mitral/tufted cells phase-lock strongly to the fast oscillations evoked by fictive odorants, and exhibit tightly coupled spike-spike synchrony on the gamma timescale during this stimulation. Moreover, the specific population of synchronized neurons differed based on the “quality”, but not the “concentration”, of the fictive odorant presented, and was conserved across multiple presentations of the same fictive odorant. Given the established selectivity of piriform cortical pyramidal neurons for inputs synchronized on this timescale, we conclude that spike synchronization on a milliseconds timescale is a metric by which the OB encodes and exports afferent odor information in a concentration-invariant manner. As a corollary, mitral/tufted cell spikes that are not organized in time may not contribute effectively to the ensemble odor representation.

## Introduction

Sensory systems are faced with the task of encoding information about an organism’s environment in an accurate yet efficient manner. At the periphery, primary sensory neurons typically represent stimuli within their receptive fields by changes in their mean spike rates that vary on the behavioral timescale in accordance with evolving stimulus properties. Subsequent stages of sensory processing, however, can transform and embed this sensory information into temporally structured activity that operates on faster timescales and governs the intra- and inter-areal synchronization properties of neuronal ensemble spike patterns. In olfaction, such fast-timescale spike synchronization is known to be necessary for fine odor discrimination in the honeybee (Stopfer, Bhagavan, Smith, & Laurent, 1997) and its importance in mammalian olfaction is becoming evident as well.

In the vertebrate olfactory bulb (OB), the representation of odor identity has long been understood to be distributed across populations of differently-tuned principal neurons (mitral and projecting tufted cells; MTCs). Odor-selective ensemble activity patterns can be identified in MTC activity rates on the behavioral timescale – specifically, in the slower periodicity associated with respiration or active sniffing behaviors (theta band; 2-12 Hz in rodents; Gervais, Buonviso, Martin, & Ravel, 2007). However, it also has long been appreciated that odor sampling generates fast oscillations in the OB local field potential (*gamma* oscillations; 30-100 Hz; Freeman, 1978), indicating that neuronal ensemble activity in the OB becomes intrinsically organized at this faster timescale in response to afferent inputs. (These OB-intrinsic oscillations then can transition into the somewhat slower *beta* band (15-30 Hz) during epochs of coherence with piriform cortex; (Frederick et al., 2016). Physiological recordings and theoretical analyses have described a dynamical system arising from the reciprocal interactions of resonant principal neurons (MTCs; Desmaisons, Vincent, & Lledo, 1999; Rubin & Cleland, 2006) and inhibitory interneurons (granule cells; GCs) within the OB external plexiform layer (EPL) (Lagier, Carleton, & Lledo, 2004; Li & Cleland, 2017; Peace et al., 2018; Schoppa & Urban, 2003). This PRING network (pyramidal resonance interneuron network gamma; Li & Cleland, 2017; Peace et al., 2018) is able to generate stable gamma oscillations that are zero-phase coherent across the EPL (Freeman, 1978; Kay et al., 2009; Kay & Lazzara, 2010), remain robust to heterogeneous MTC activation levels (Cenier et al., 2008; Li & Cleland, 2017; Lowry & Kay, 2007) and constrain the timing of MTC spikes to a particular gamma phase window (Bathellier, Lagier, Faure, & Lledo, 2006; Kashiwadani, Sasaki, Uchida, & Mori, 1999). More specifically, the dynamical system generated by this cellular network temporally phase-constrains the spiking of activated MTCs with respect to its coherent periodicity on the gamma timescale, thereby generating an oscillatory local field potential as a side effect of this coordinated activity. However, the functional role of this gamma-timescale temporal structure in odor representations – specifically, the extent to which MTCs encode odorant quality information in their spike timing properties on the gamma timescale – remains unknown.

These limitations on our understanding persist in part because odorant stimuli are difficult to control. All odorants – even monomolecular odorants – bind to multiple types of primary odorant receptors with different affinities and efficacies (discussed in Gronowitz, Liu, Qiu, Yu, & Cleland, 2021), such that primary olfactory sensory neuron (OSN) activation profiles vary nonlinearly and sometimes non-monotonically with concentration. Moreover, the timing of odor stimulus delivery is imprecise, rendering it difficult to determine whether variations in fine-scale temporal response properties arise from internally regulated OB network dynamics or simply from the undercontrolled timing of odorant delivery to receptors. We here use an *in vitro* olfactory bulb slice preparation to overcome these constraints. Specifically, we here deliver dynamically “odor-like” stimuli to OB slices using spatiotemporally patterned optical stimulation of OSN axonal arbors in OB slices, while recording MTC ensemble responses using a 120-channel planar multielectrode array. This strategy enables a mechanistic assessment of stimulus-evoked fast oscillatory dynamics in the EPL network, and illustrates how the physical features of sensory inputs are reflected in MTC response profiles, including the regulation and synchronization of MTC spike times on the gamma timescale.

The synchronization of convergent action potentials on fast (gamma/beta) timescales is well-matched to the typical integration time constants of neuronal membranes, and thereby enables effective heterosynaptic integration in follower cells. In particular, it is established that pyramidal neurons in the piriform cortex are selectively activated by convergent MTC spike inputs that are synchronous on this timescale (Luna & Schoppa, 2008). This selectivity suggests that MTCs would effectively represent odorants via patterns of spike synchronization on this gamma/beta timescale. We therefore sought to ask: how do odor-evoked dynamics intrinsic to the OB shape MTC spiking activity? Are the resulting patterns of spike synchronization on the gamma timescale both *replicable* and *odor-specific?* And, finally, in addition to this metric transformation of the glomerular-layer representation into a fast spike synchronization-based representation on the gamma timescale, does a computational transformation of information content also occur?

## Results

### Optical stimulation with “fictive odors”

Horizontal OB slices were taken from OMP-ChR2/EYFP (ORC-M) mice and placed onto a 120-channel planar multielectrode array (MEA). Each slice was imaged with a custom fluorescent microscope (see *Methods*) to determine the location of the EYFP-expressing glomerular layer within the slice (Figure 1A). Spiking activity was observed along a band of electrodes located immediately deep to the glomerular layer (corresponding to the external plexiform and mitral cell layers), and could be evoked by optical stimulation of the glomerular layer (Figure 1B-C).

**Figure 1.**
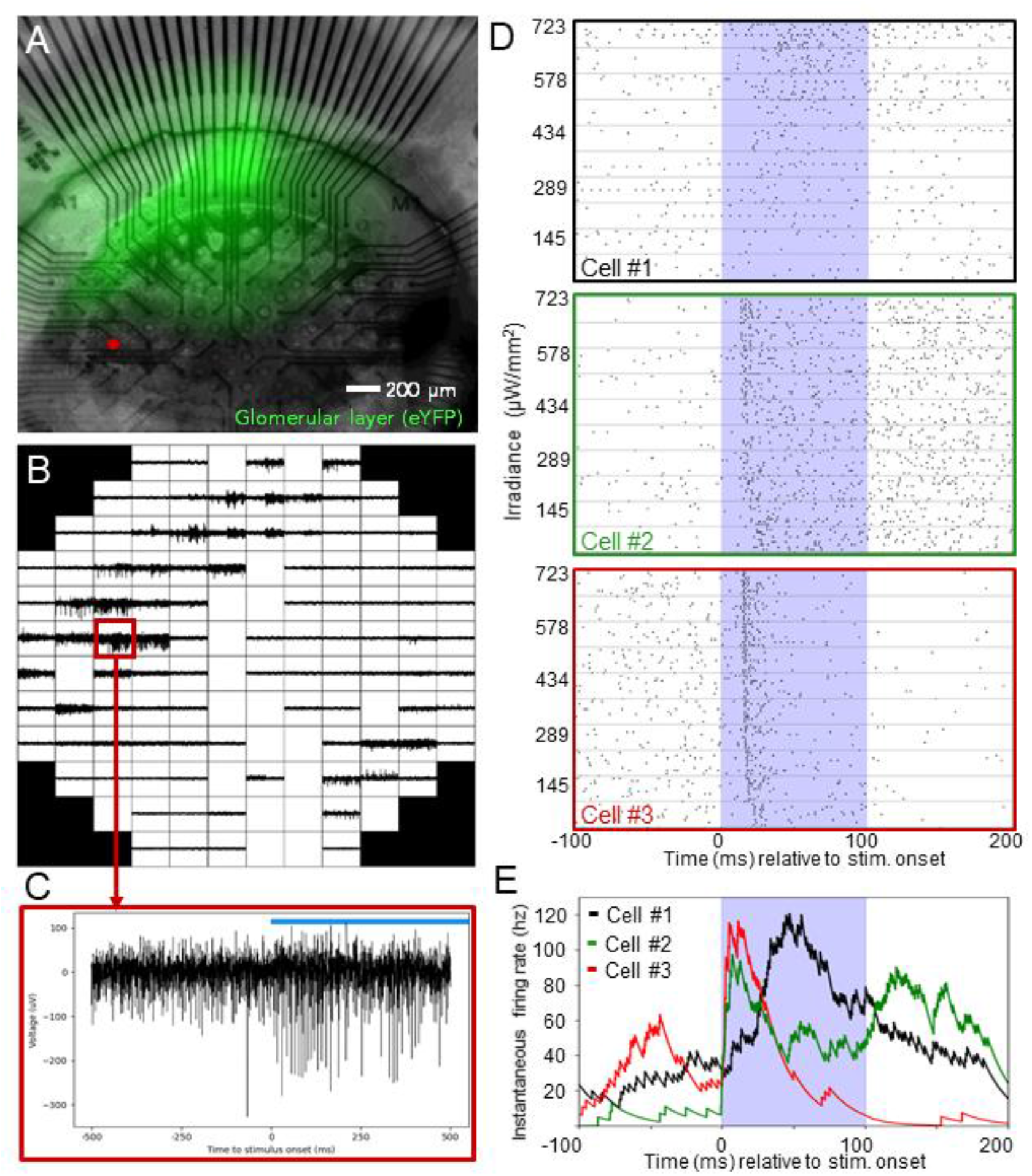
Bulbar responses to optical stimulation. **(A)** Horizontal OB slice with fluorescing glomerular layer (green) shown on top of a 120-electrode array (50% transparency overlay). The location of the electrode shown in panel C is indicated (red dot). **(B)** 120-channel MEA recording showing a band of increased LFP power and spiking activity corresponding to the EPL/MCL layers in the OB slice. **(C)** Single electrode recording showing spiking activity from presumed MTCs evoked by optical stimulation (blue bar). **(D)** Raster plots of MTC responses to a series of 100 ms light pulses presented at increasing light intensities. Individual rows within each raster each contain 10 trials delivered at a single intensity. **(E)** Instantaneous firing rates of the cells shown in panel D in response to light stimulation. Spike trains were convolved with a negative exponential kernel (see Methods) and the results averaged across the ten trials with the highest stimulus intensity (723 μW/mm^2^).

The parameters of light stimulation were designed to provide naturalistic, odor-like afferent input to the slice. First, we calibrated stimulus intensity by characterizing neural responses to a single pulse of light across a series of increasing intensities. We determined a range of light intensities (0.1-1.0 uW/mm^2^) that drove robust increases in the instantaneous firing rates of MTCs, comparable to odor-evoked firing rates and patterns recorded *in vivo* (Figure 1D). Moreover, within this intensity range, for any given optical stimulus pattern, the relationship between light intensity and the stimulus-evoked firing rate varied across responsive MTCs – exhibiting a response diversity comparable to the distinct classes of responses described in foundational studies of MTC odorant responses (Hamilton & Kauer, 1989; Meredith, 1986; Wellis, Scott, & Harrison, 1989). Many neurons showed stimulus-evoked increases in firing rate that either returned to baseline shortly after the stimulus ended or were sustained well beyond the stimulus duration; other neurons, in contrast, showed periods of clear suppression subsequent to their initial excitation (corresponding respectively to E1 and E2 - type responses in Hamilton & Kauer, 1989; Meredith, 1986; Figure 1D-E). Wholly suppressive responses (S-type) were observed relatively rarely (see Figure 2), in part because of the lower baseline MTC spike rates observed *in vitro*, and perhaps also because the slicing process reduces lateral connectivity in the deep glomerular layer, which normally contributes to feedback normalization (Banerjee et al., 2015; Cleland & Borthakur, 2020; Peace et al., 2018).

**Figure 2.**
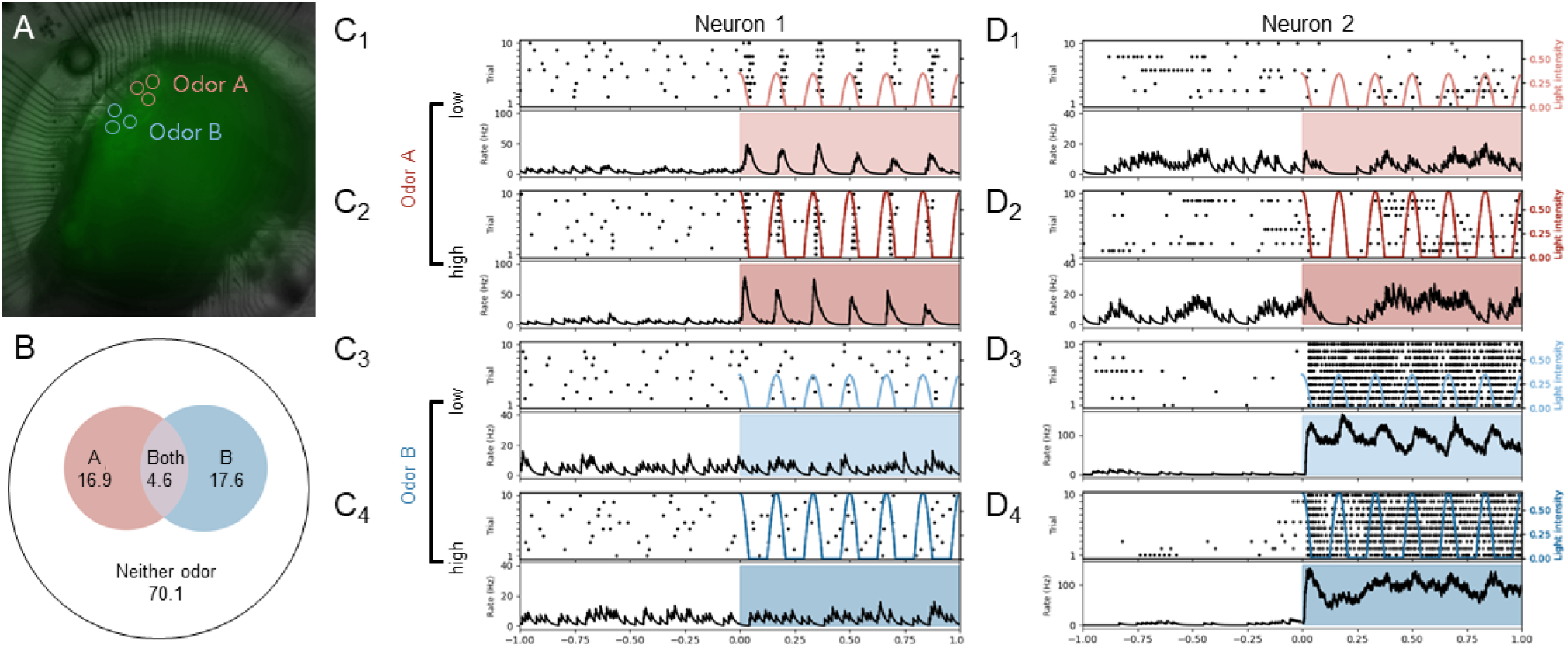
MTC responses reflect fictive odor quality. **(A)** The regions of stimulation comprising two fictive odors (“A” and “B”) on an OB slice. **(B)** Diagram depicting the excitatory responses to fictive odors A and B as a percentage of all MTCs recorded in the slice. **(C)** Responses of an individual neuron to stimulation with fictive odors A and B, each presented at low (250 uW/mm^2^) and high (500 uW/mm^2^) peak intensities. *Top frame:* raster plot of 10 responses to fictive odor delivery, overlaid with the timecourse of the periodic light stimulus. *Bottom frame:* Instantaneous firing rate averaged across trials. Highlighted regions indicate stimulus duration. **(D)** Same as C1-C4, with a different MTC, selected to highlight response variability.

To model the sparse glomerular activation characteristic of natural odorants, each spatiotemporal stimulus pattern (“fictive odor”) consisted of three localized spots of light (60 μm radius), delivered onto the glomerular layer with 6 Hz (theta band) intensity modulation to reflect active sniffing dynamics. Within a given slice, we defined 2-4 stimulus patterns that differed in “quality” (i.e., that differed in which glomeruli were stimulated; see *Stimulus Modeling* section in *Methods;* Figure 2A). In some experiments, each fictive odor stimulus was presented at different “concentrations’’, wherein the intensities of light-activated regions were adjusted co-monotonically within the established intensity range. Under these stimulation parameters, 5-35% of the MTCs recorded in each slice showed excitatory responses to a given fictive odor (Figure 2B); the remainder exhibited no response, or else exhibited suppressive/inhibitory responses that could not be clearly distinguished given the low baseline MTC spike rates. Fictive odors that differed in quality evoked excitatory responses in different ensembles of MTCs, and these ensembles of activated MTCs were largely conserved over different intensities (“concentrations”) of the same fictive odor (Figure 2C-D).

### Stimulus-evoked gamma rhythmicity in the local field potential

Fictive odor stimuli evoked persistent gamma oscillations in the LFP (Figure 3A-B), consistent with previous results from our lab (Peace et al., 2018) and with natural responses to odors recorded *in vivo* (Freeman, 1978; Kay et al., 2009). In OB slices, electrodes with measurable gamma oscillations formed a band immediately deep to the glomerular layer, consistent with the location of the external plexiform and mitral cell layers (Figure 1A-B; Figure S1). Stimulus-evoked gamma oscillations commonly occurred in an odor quality-specific manner, such that power levels in the gamma band measured at multiple recording sites covaried across different intensities of a particular fictive odor (see *Methods*), but varied relative to one another in response to fictive odors of different qualities (Figure 3A-B).

**Figure 3.**
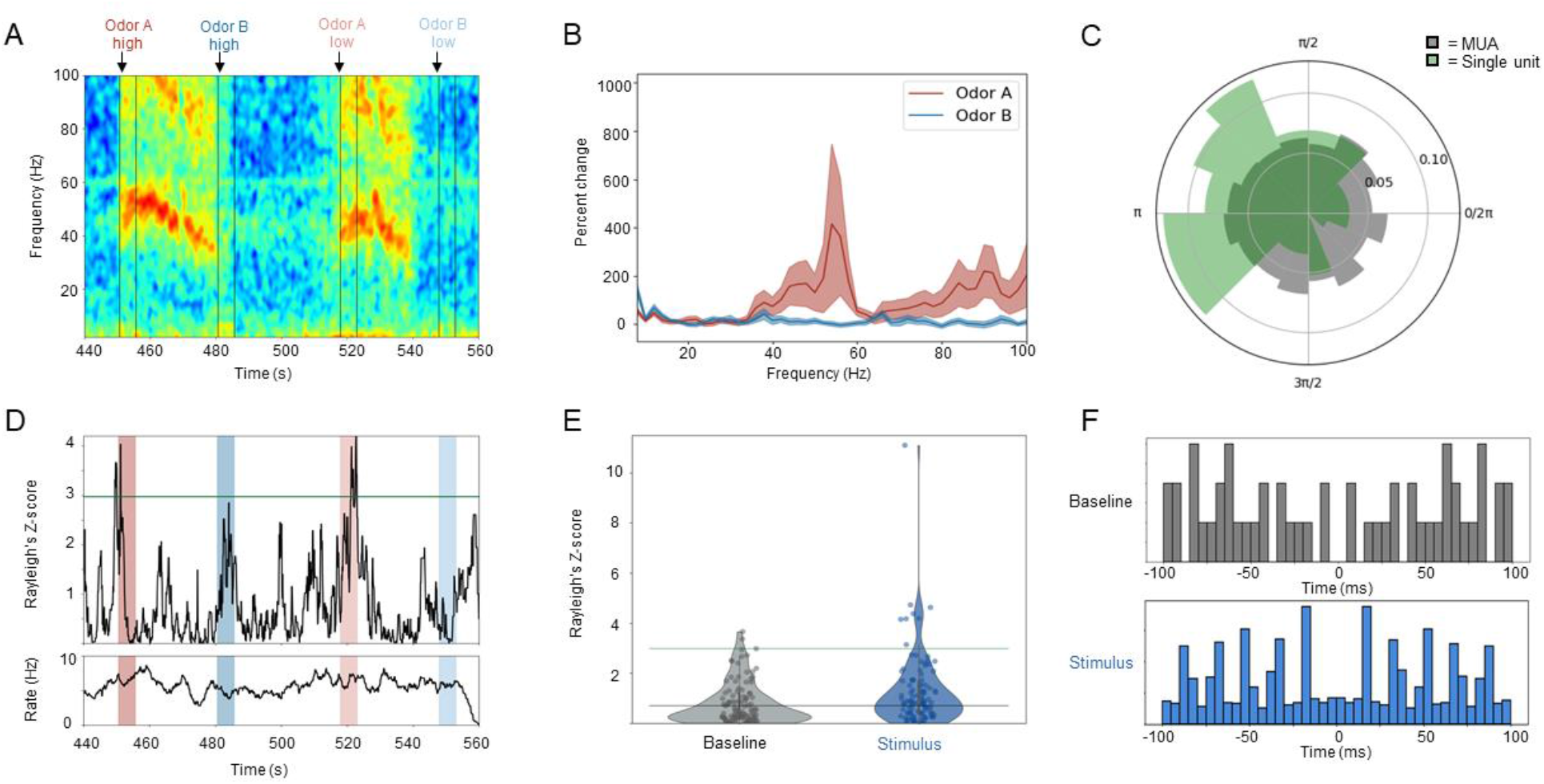
Gamma rhythmicity in MTC responses to fictive odor presentation. **(A)** Spectrogram showing stimulus-evoked gamma oscillations recorded from one selected electrode in response to fictive odors A and B (as depicted in Figure 2A), each presented for five seconds (delineated by vertical lines) at high and low intensities (same as in figure 2). Evoked gamma oscillations persisted following stimulus termination. **(B)** Power spectra from the same electrode as panel A, measured during the five-second stimulus presentation and averaged across 20 trials for each odor (low and high intensities combined). **(C)** Circular histogram illustrating the spike phase distribution for all recorded MTCs combined (gray) and for a selected individual MTC (green) during stimulus-evoked gamma oscillations. In both cases, the distributions have a non-uniform distribution of phases (*p*<0.05 and *p*<0.001 for combined multi-unit activity (MUA) and single-unit responses, respectively). **(D)** *Top:* Time-resolved (5-second window, 100-ms step size) Z-score for the phase-locking of an individual neuron to the LFP gamma band; the green line indicates the Z-score corresponding to a *p*-value of 0.05. *Bottom:* Instantaneous firing rate of the same neuron (same window and step size). Both plots are from the same recording, and share the same time axis, as the spectrogram in panel A. Highlighted windows indicate periods of fictive odor stimulation. **(E)** Distribution of Rayleigh’s Z-scores for phase-locking of the MTC population in a single slice (*n* = 104 single units) during the pre-stimulus baseline (*gray*) and high intensity fictive odor stimulus (*blue*) periods. Horizontal lines denote *p* = 0.5 (bottom/gray) and *p* = 0.05 (top/green). **(F)** Autocorrelograms for unit activity during baseline (*gray*) and stimulus (*blue*) periods.

Oscillations in the LFP reflect temporal coordination among neural assemblies in the local population. Dynamical systems-based coordination of the membrane properties of neurons in such assemblies can robustly constrain the timing of action potential generation and propagation (Li & Cleland, 2017). Therefore, we first sought to determine whether spiking activity in recorded MTCs was phase-constrained with respect to fictive odor-evoked gamma oscillations by computing the phase-locking values (PLV) of unit activity – that is, the phase-specificity of evoked action potentials with respect to simultaneously recorded LFP gamma oscillations (see *Methods*). Because the PLV measure is biased with respect to spike count, we subsequently applied Rayleigh’s uniformity transformation, generating Z-scores and corresponding *p*-values for each PLV measure that compensate for this bias. Unsorted multiunit activity demonstrated weak but significant phase-locking to stimulus-evoked gamma oscillations (*p* < 0.05, Rayleigh’s test of uniformity; 167.8° phase preference; Figure 3C, *MUA*), whereas strong phase-locking was clearly present in the activity of a subpopulation of individual neurons (*p* < 0.01, 136° phase preference for unit depicted;Figure 3C, *Single unit*). This result suggested that fictive odor stimulation might selectively constrain spike phases in a stimulus-specific population of MTCs. Time-resolved analysis revealed that the gamma phase-constraint of action potentials generated by individual stimulus-activated MTCs indeed increased specifically and transiently during stimulus presentation. Notably, this stimulus-gated phase constraint could occur even in units in which the stimulus did not increase the instantaneous spiking rate (Figure 3D).

In principle, single-unit spike trains with non-Poisson statistics could generate phase distributions over a time window of finite length that systematically differ from Rayleigh’s null hypothesis of uniformity, even in the absence of underlying periodicity. For example, a rapid burst of activity from a neuron could yield several spikes that are similar in phase yet unrelated to any coordinating network oscillation. To determine the extent to which activity in a recorded population of neurons would be likely to generate non-uniform phase distributions without any periodic modulation of spike timing, we assessed the phase-locking of single unit activity in the pre-stimulus period (during which there was no detectable spectral peak in the gamma band). This measure then was used as a baseline against which to compare the properties of stimulus-evoked PLVs. On average, fictive odor stimulation significantly increased the gamma-band phase-locking of individual MTCs over that measured at baseline (Rayleigh Z-scores of PLVs; Z_PLV_ = 1.276 compared to baseline Z_PLV_ = 0.817; paired *t*-test; *t*(103) = 2.85, *p* < 0.01; Figure 3E). Moreover, analyses of individual units revealed considerable variability among MTC responses. Specifically, whereas spiking activity in most MTCs appeared relatively unaffected by the presentation of a particular fictive odor, a substantial minority was strongly phase-constrained to gamma (7.7% with *p* < 0.05, compared to 1.9% at baseline; Figure 3E). Gamma rhythmicity also was clearly present in the autocorrelograms of MTC units during the stimulus period, whereas the pre-stimulus baselines lacked any obvious rhythmicity (Figure 3F). Together, these results indicate that fictive odor stimulation – i.e., selective glomerular activation via delivery of specific spatiotemporal patterns of light – drives fast, rhythmic responses in the OB external plexiform layer that phase-constrain the action potentials of selected individual MTCs in the gamma band.

### Spike synchronization-based representations of fictive odorants

Based on this elevated phase-locking of MTC spikes to LFP gamma, we hypothesized that fictive odor stimulation increased the gamma-timescale synchronization of spikes generated by co-activated MTCs. To assess spike-spike synchronization, we conducted unitary event (UE) analyses (Grun, 2009; Grun et al., 2002a; see Methods) on spike trains from all pairwise combinations of recorded MTCs during fictive odor presentation. Briefly, this method is designed to detect coincident spike events between parallel spike trains on a defined timescale. We used a bin size of 5 ms to reflect the *permissive epoch* (Imam & Cleland, 2020) of a gamma cycle – i.e., roughly 25% of a 50 Hz oscillatory period. The significance of coincident spiking was evaluated by generating a distribution of expected coincidences for each pair of spike trains based on bootstrapped surrogate data in which spike times from the real data were randomly dithered by up to 10 ms. Critically, this surrogate approach conserves key statistical properties of the original data – such as the instantaneous firing rate and approximate spike train structure – while destroying the fine temporal precision of spike events. Unit pairs were deemed “synchronous” only if the number of their empirical spike coincidences was in the top 1% of their respective surrogate distributions.

A representative example of UE analysis on a pair of units activated by fictive odor stimulation is shown in Figure 4A. Synchronized spiking (within a 5 ms bin) increased during the stimulus window, was usually modulated by the phase of the periodic stimulus input (6 Hz *theta* band, mimicking mouse sniffing behavior), and remained elevated throughout the stimulus presentation even as the firing rates of the units returned to their pre-stimulus baselines. Because of the considerable diversity in synchrony responses among MTC pairs, additional examples of UE analyses are presented in supplementary figure S2. At baseline, only 1.2% of MTC unit pairs showed synchronous spiking activity, indistinguishable from chance levels (i.e., 1%). During fictive odor stimulation (1.0 seconds immediately following stimulus onset), 2.3% of all unit pairs fired synchronously, significantly more than baseline levels (two-sample *t* test, *p* < 0.001, Figure 4B).

**Figure 4.**
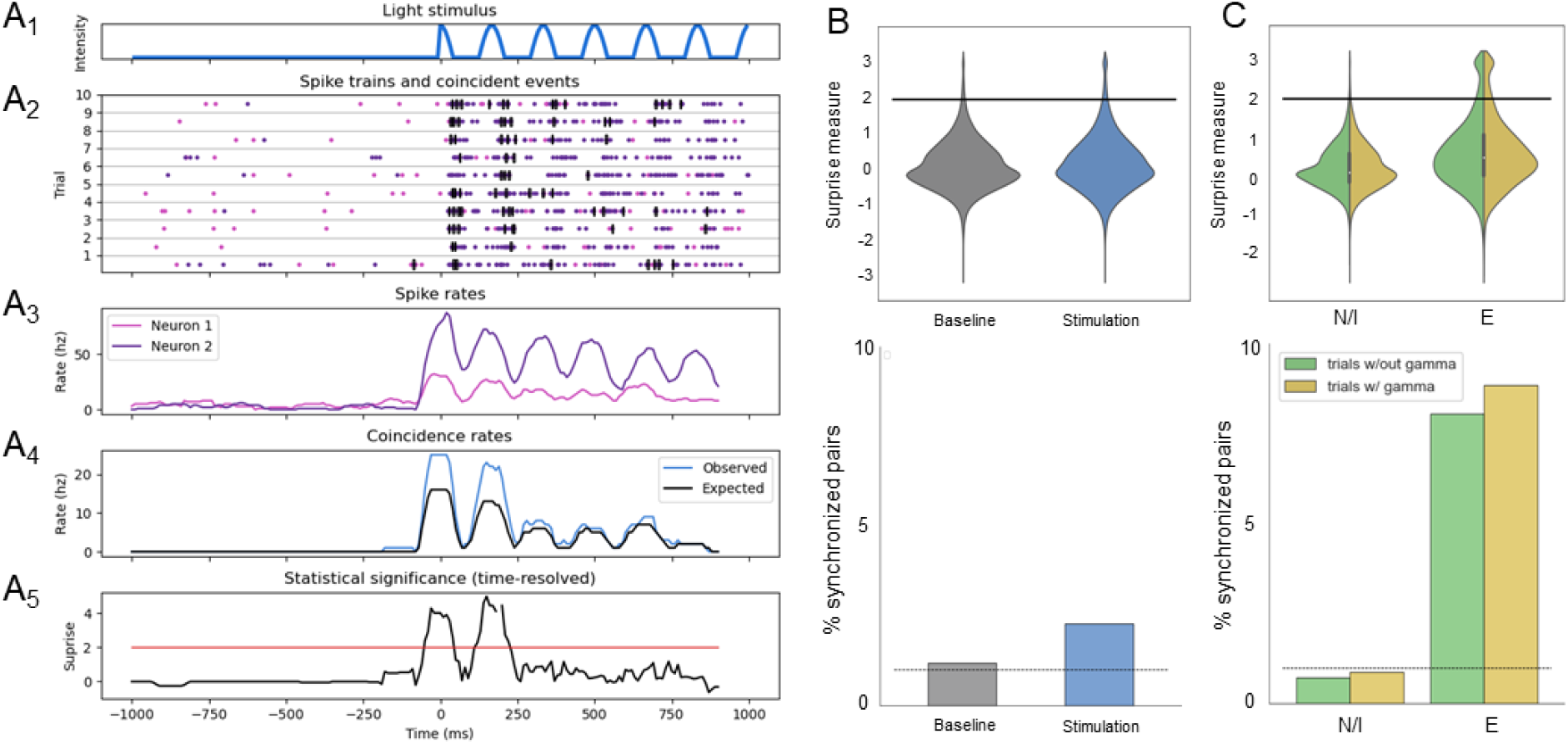
Fictive odor stimulation increases MTC-MTC pairwise synchrony. **(A1)** Light intensity profile during a single fictive odor trial. **(A2)** Raster plots for two MTCs (*purple* and *pink*) in response to stimulation; each row is a single trial aligned to stimulus onset. Bins with coincident events are indicated with black rectangles. **(A3)** Instantaneous firing rates for the same neurons shown in panel B. **(A4)** Observed rates of coincident spiking between these two neurons, and expected coincident rates based on the surrogate data. **(A5)** Time-resolved statistical significance (logarithmically plotted as the surprise measure) of the coincidence rate. Red line at surprise = 2.0 indicates a significant level of spike synchrony at *p* = 0.01. **(B)** Distribution of surprise values (*top*), and percentages of pairs that are synchronized (*bottom*) for all recorded MTC units at baseline and during fictive odor stimulation (*n* = 625 neurons, *N* = 5 slices). Horizontal bar in upper panel denotes significance boundary (α = 0.01); bar in lower panel indicates the percentage of synchronized pairs expected by chance (1%, based on α = 0.01). **(C)** Distribution of surprise values (*top*), and percentages of synchronized pairs (*bottom*) for MTC units based on their responses to fictive odor stimulation (*N//*: non-responsive/inhibitory, *E:* excitatory), and on whether visible gamma rhythmicity was evoked on the reference electrode on a given trial (*n* = 209 units with E-type responses, 557 units with N-type responses, *N* = 5 slices).

Next, we asked whether fictive odor-evoked synchrony varied among MTCs based on how they responded to stimulation. Fast excitatory responses were easily identifiable in our data; however, at the low spontaneous firing rates commonly observed in *in vitro*, inhibitory responses could not easily be discerned from nonresponsiveness, and quantitatively screening for inhibition proved impractical. As a result, we dichotomously classified MTC responses to any given fictive odor as either “excitatory” or “non-responsive/inhibitory” (see Methods). Whereas only 0.8% of unit pairs fired synchronously when neither unit displayed an excitatory response – a rate indistinguishable from chance – 8.4% of pairs of units with excitatory responses were synchronized (Figure 4C; see below for analysis).

We next sought to determine whether the presence of measurable LFP gamma oscillations corresponded to increased rates of gamma-timescale spike synchrony in the recorded population. To test this, we identified the electrode with the strongest stimulus-evoked changes in gamma power, and grouped trials based on whether or not gamma was present on this electrode – i.e., whether overall EPL network activity was sufficiently strong and coordinated to generate a clear oscillatory field potential on the gamma timescale. Interestingly, we found that synchrony was modulated by gamma in a “response type”-dependent manner. Specifically, 8.1% of unit pairs that displayed excitatory responses showed single-trial synchrony when gamma was not measurable, whereas 8.8% of excitatory unit pairs were synchronized on trials that also evoked clear fictive odor-evoked gamma. Unit pairs in which both units displayed non-excitatory responses exhibited single-trial synchrony in 0.72% of trials in the absence of detectable gamma rhythmicity, compared to 0.87% of trials with clear gamma. Two-factor ANOVA indicated significant effects of both response type (F(1,103306) = 3571, *p* = 1.11e-16) and the presence of visible LFP gamma (F(1,103306) = 4.28, *p* = 0.038) on synchrony among pairs of MTCs. The interaction was not significant (response type*gamma: F(1,103306) = 1.37, *p* = 0.24). These results suggest that while afferent stimulation can evoke spike synchronization without increasing MTC spike rates (Figure 3D; supplementary Figure S2), it is more common that synchronized spikes arise from excited MTCs. However, the majority of MTCs, excited or not, do not synchronize their action potentials on the gamma timescale in response to any given afferent stimulation. MTC spike synchronization is a selective phenomenon.

Fine timescale synchrony among pairs of MTCs varied across trials in accordance with the stimulus presented (Figure 5A-B). To systematically characterize the dependence of MTC synchrony on the quality and concentration of the fictive odor stimuli, we constructed response synchrony matrices for each trial composed of the surprise values between each pair of neurons recorded in the slice. These high-dimensional matrices then were projected onto a three-dimensional principal component (PCA) space for visualization (Figure 5C). We used distance in PCA space to quantify the difference in response synchronization patterns to fictive odor stimuli of different qualities and intensities (“concentrations”); each distance was normalized to the mean distance between repeated presentations at the same quality and intensity. The synchronization patterns evoked by fictive odors of different quality were significantly more different from one another than responses on repeated trials (one-way ANOVA, F(2, 957) = 39.4, *p* =- 4.91e-17, followed by Tukey HSD post hoc test, *p* < 0.001, Figure 5D), indicating that inter-odor variance in synchrony responses is greater than intra-odor variance, and that fine-scale patterns of spike synchrony are conserved in odor representations and can distinguish odorants of different qualities from one another. In contrast, the synchronization patterns evoked by a given fictive odor at different “concentrations” showed no greater difference than repeated trials at the same intensity (Tukey HSD; *p* = 0.90; Figure 5D), indicating substantial concentration invariance in the spike synchronization pattern of odor responses. Combined with the established selectivity of piriform cortex (PCx) for synchronized inputs (Luna & Schoppa, 2008), this finding suggests that the improved concentration invariance of PCx principal neuron responses compared with bulbar MTC responses (Bolding & Franks, 2018) arises from MTC synchronization properties That is, PCx neurons are selective for a feature of MTC responses – fast timescale spike synchronization – that discards the residual concentration-dependent stimulus variance observed in MTC responses when assessed at slower timescales. This selectivity then would conclude a multistage procedure for distinguishing stimulus variance arising from odorant quality from that arising from concentration differences (Cleland et al., 2011).

**Figure 5.**
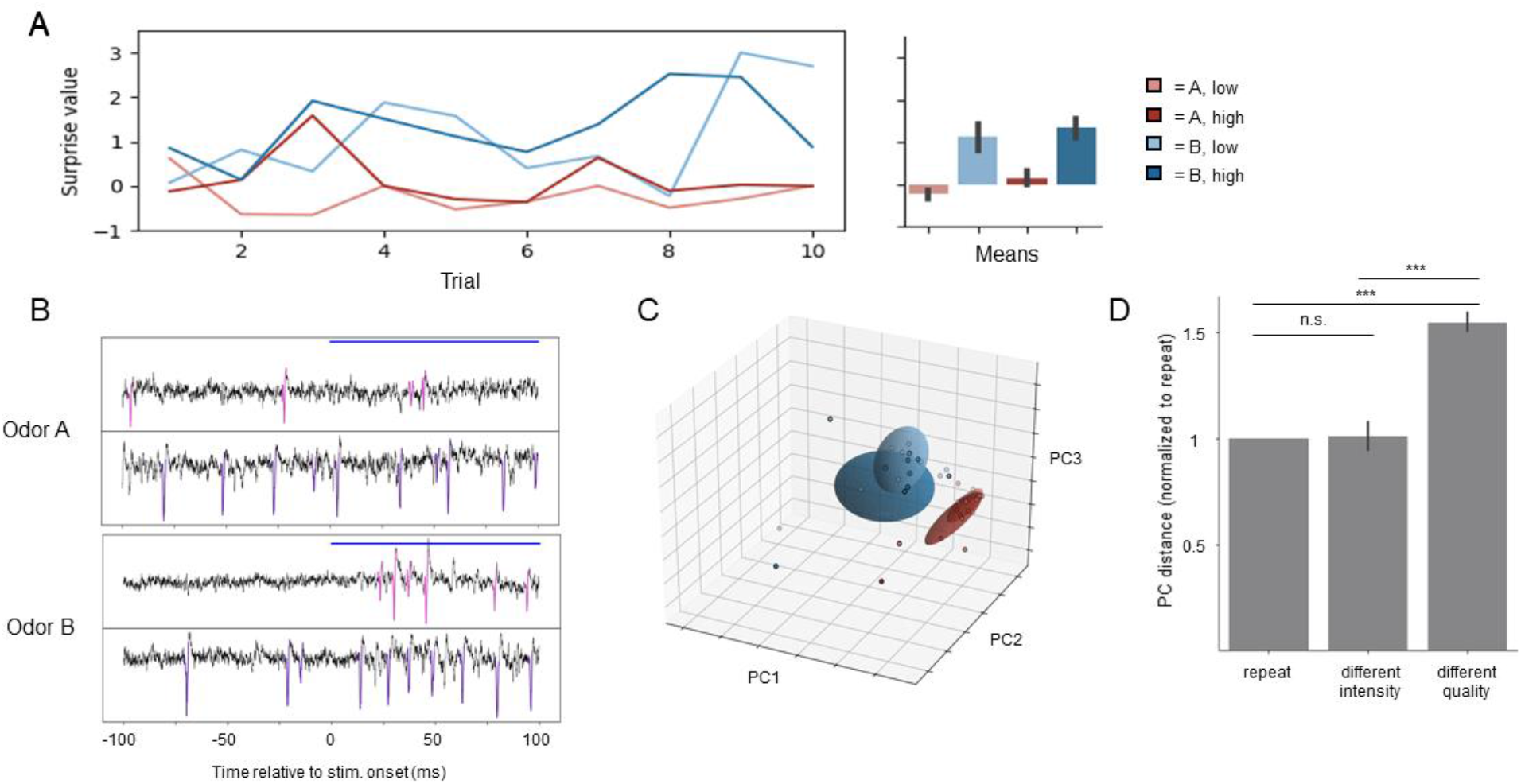
Quality-specific spike synchrony. **(A)** *Left:* Surprise values of pairwise spike synchronization from an example pair of MTCs across 10 fictive odor trials. Spikes from this MTC pair are synchronized in response to fictive odor B (at both intensities), but not to fictive odor A. *Right:* Surprise values averaged across 10 trials for each condition. **(B)** Single trials of fictive odors A and B (high intensity) for the same pair of neurons as in panel A. Spikes are colored in pink (neuron 1, *upper panels*) or purple (neuron 2, *lower panels*). **(C)** Principal component analysis (PCA) representation of the synchrony matrix for all unit pairs in a single slice in response to the presentation of fictive odors A (red) and B (blue) at different “concentrations” (color saturation denotes relative intensity). Each dot represents an individual trial, and ellipsoids denote the mean ± 1 SD for each condition. **(D)** Average pairwise distances between repeated trials of a fictive odor stimulus at the same quality and concentration, trials that differed only in “concentration”, or trials that differed in quality, all normalized to the average intra-stimulus (*repeat*) distance. Changes in fictive odor quality generate different synchrony matrices (*p* <.001), whereas changes in intensity do not (*p* = 0.90).

## Discussion

Spike synchronization is important for information processing and interareal communication across the brain. In the olfactory system, odors evoke synchronized spiking among MTCs and fast oscillations in the bulbar LFP, but the study of how olfactory information might be organized within a fast temporal framework has been challenging because of experimental limitations such as the low temporal resolution and replicability of odor stimulus delivery. In the present study, we used optogenetics to present spatiotemporally patterned stimuli to OB slices while recording MTC ensemble responses electrophysiologically – an approach which enabled the presentation of “odor-like” stimuli with a level of statistical precision and replicability not possible with real odorants. These fictive odors evoked responses comparable to natural odor-evoked responses in terms of MTC unit firing rates, classical MTC response types, and fast LFP oscillations. Importantly, MTC responses also were marked by synchronized spiking – measured both as phase-locking to stimulus-evoked gamma oscillations and as patterns of increased spike synchrony among unit pairs. Spiking activity was most strongly synchronized among pairs of units with fast excitatory responses, and was modestly more synchronous on trials where the presented fictive odor evoked measurable gamma rhythmicity in the LFP. To dissect how the spike-timing properties of MTCs varied with odor “quality” and “concentration”, we systematically varied the parameters of fictive odor stimuli based on explicit models of these natural odor properties. We found that fast-timescale spike synchronization patterns were diagnostic of the “quality” of the fictive odor presented, and were conserved when the same fictive odor was presented at different “concentrations”. Specifically, we computed *N*x*N* synchrony matrices consisting of pairwise surprise values among *N* recorded units, such that stimulus responses mapped to different manifolds in the matrix as a function of the quality of the fictive odor stimulus presented (Figure 5C).

This analytical approach was based on a pairwise measure that reliably captured synchrony between spike trains in the data, which then was generalized to all possible unit pairs to generate ensemble response data. In principle, a nonpairwise metric of synchrony (such as the PLV) could have been used instead to construct an *N*-dimensional vector of each unit’s synchronization with overall population activity, as could be assessed by LFP recordings of gamma oscillations. We favored the pairwise unitary event analysis approach, because it is spike-spike synchrony, and not LFP gamma *per se*, that exerts downstream computational effects, and also because spike-LFP measures are less reliable as a method for comparing synchronization patterns across stimulus conditions. To wit, reliable LFPs were not always detected on recording electrodes for the diverse fictive odors presented, and could vary considerably across trials.

Coding schemes based on spike timing provide considerable advantages over those based solely on firing rates. For example, encoding information in firing rates is slow, because the outcome cannot be interpreted until the integration window over which the mean rate can be reliably calculated has concluded. In contrast, the activity of an ensemble of synchronized MTCs can be reliably communicated within a single gamma/beta cycle, and the analogue activation levels of neurons in a synchronized ensemble can be communicated unambiguously via spike phase (relative latency) (Linster & Cleland, 2010). Recent olfactory bulb models have used such gamma-modulated spike timing-based representations together with iterative attractor dynamics to solve a critically underappreciated problem in olfaction: how to recognize known odors despite the presence of destructive interference from competing background odors, in addition to the natural variance resulting from odor plume dynamics (Imam & Cleland, 2020).

### Concentration invariance arises from spike synchronization

The olfactory system samples odor inputs that vary in concentration by orders of magnitude, yet the perceptual identity of an odor is largely stable across broad concentration ranges (Homma et al., 2019). This is a nontrivial problem, as odor quality and concentration both are mediated by changes in the activation profiles of odorant receptors that are not fully dissociable (see *Methods: Stimulus modeling*). The early olfactory system deploys a series of concerted mechanisms to segregate quality variance from concentration variance (Cleland et al., 2012); however, even at the level of bulbar MTCs, odor responses still vary considerably with odor concentration (Bolding & Franks, 2018; Hamilton & Kauer, 1989; Imamura, Mataga, & Mori, 1992). Odor representations in PCx, by comparison, are much more concentration-invariant, which has led to the hypothesis that PCx selectively extracts the most concentration-invariant feature(s) of its OB inputs (Bolding & Franks, 2018). Owing to the apparent concentration invariance of MTC odor representations when framed within a gamma-timescale spike synchronization metric (Figure 5), and the selectivity of PCx neurons for MTC spikes synchronized on that timescale (Luna & Schoppa, 2008), we propose that *fast-timescale spike synchronization* is that concentration-invariant feature. Notably, higher stimulus concentrations still may generate tighter synchrony among the ensemble of activated MTCs, thereby affording a means of communicating stimulus intensity so as to represent, for example, the intensive salience of an odor (Cleland & Borthakur, 2020; Cleland, Narla, & Boudadi, 2009).

The odor-specificity of patterns of fast-timescale spike synchrony among MTCs – and the conservation of such patterns across stimulus intensity levels – indicate that these synchronization patterns can provide an effective physiological basis for the OB to communicate odor information to PCx and other projection targets. In this framework, gamma and beta field potential oscillations reveal the underlying dynamical organization of the olfactory system’s cellular networks, organizing the timing of stimulus-relevant MTC spikes for effective interareal communication. Importantly, spikes emitted from active MTCs that are *not* synchronized to these oscillations would not effectively influence the activity of follower cells, whereas less active MTCs could still be recruited into active ensembles by careful reorganization of their spikes in time. The importance of MTC spike timing regulation affords granule cells, and by extension, the PCx feedback fibers which innervate them, a powerful and subtle influence on the nature of the information that MTCs export from the OB.

## Methods

### Transgenic mice

Olfactory bulb slices were prepared from OMP-ChR2/EYFP (ORC-M) plasmid transgenic mice of both sexes (Dhawale, Hagiwara, Bhalla, Murthy, & Albeanu, 2010), initially provided by Venkatesh Murthy, Harvard University. These mice coexpress channelrhodopsin-2 (ChR2) and enhanced yellow fluorescent protein (EYFP) under the control of the olfactory marker protein (OMP) promoter, thereby targeting transgene expression to all OSN axonal arbors within the OB glomerular layer. All procedures for MEA experiments were performed under the auspices of a protocol approved by the Cornell University Institutional Animal Care and Use Committee (IACUC). Cornell University is accredited by the Association for Assessment and Accreditation of Laboratory Animal Care (AAALAC International).

### Microscopy and stimulus delivery

Optical stimulation was delivered by a high-speed (up to 1.44 kHz refresh rate) digital light processing (DLP) projector (PROPixx, VPixx Technologies) utilizing RGB LEDs and a digital micromirror device (DMD) for stimulus control, with the stimulus image focused onto the OB slice by a customized series of lenses (Thorlabs Cerna microscope system; *https://cplab.science/meascope*). Images of the OB glomerular layer were taken by briefly illuminating the slice with 500±10 nm green light (Figure 1A; Thorlabs FB500-10), so as to activate EYFP while not strongly activating ChR2. Using custom Ceed software (*https://cplab.science/ceed*), spatially delimited stimulus fields (e.g., Figure 2A) were defined within the glomerular layer and separately programmed with temporal profiles of stimulus intensity. The resulting spatiotemporal profiles of blue light (470±50 nm bandpass; Semrock FF02-470/100-25) then were delivered to these stimulus fields to activate ChR2-expressing OSN arbors. These “fictive odor” stimuli were presented for 5 seconds, intensity-modulated with a 6 Hz sinusoid (see Results), with intertrial intervals (ITIs) of 25 seconds.

### Stimulus modeling

How should odor stimuli be modeled using spatiotemporal patterns of light, combining what we know about the form and dynamics of natural odor activation of OB circuitry with the tighter control and replicability of optogenetic stimulation? We here take a “naturalistic” approach to this problem, posing specific hypotheses about the properties of natural odor stimulation and constructing stimuli accordingly. For example, changes in either odor quality or concentration both alter the distribution of glomerular activation levels. Concentration increases, however, generally evoke monotonic increases in the activation levels of individual glomeruli (and also may progressively recruit some new, lower-affinity glomeruli). These increases will not be linearly proportional to one another across glomeruli, but to a substantial extent the ordinal ranking of glomerular activation levels should be preserved (Araneda, Kini, & Firestein, 2000; Gronowitz et al., 2021). Quality differences, in contrast, involve sharply different ordinal rankings of glomerular activation levels. These distinctions already have been exploited in network theories of concentration tolerance (Banerjee et al., 2015; Cleland & Borthakur, 2020; Cleland, Johnson, Leon, & Linster, 2007); the present work employs them to highlight the manifestation of concentration tolerance in the patterned spiking output of MTCs.

### Slice preparation

Horizontal slices (300 μm) were prepared from the olfactory bulbs of 5-12 week old ORC-M mice. Mice were anesthetized with 2% isoflurane, injected with ketamine (150 mg/kg ip) as a neuroprotectant, and then decapitated, after which the olfactory bulbs were quickly removed. Slices were cut on a vibrating microtome (Leica VT1000S) in an ice cold, oxygenated (with carbogen: 95% O2, 5% CO2), low-calcium/high-magnesium artificial cerebrospinal fluid (aCSF) dissection solution containing (in mM): NaCl 124, KCl 2.54, NaHCO3 26, NaH2PO4 1.23, CaCl2 1, MgSO4 3, glucose 10 (Balu & Strowbridge, 2007). Slices then were incubated in oxygenated dissection solution at 37°C for twenty minutes, and then removed from incubation and maintained in this solution at room temperature until transfer to the recording well.

### Slice electrophysiology

Slices were placed on a 120-electrode planar multielectrode array (MEA; Multichannel Systems) and continuously superfused with heated (37°C), oxygenated recording aCSF containing (in mM): NaCl 125, KCl 3, NaHCO3 25, NaH2PO4 1.25, CaCl2 2, MgCl2 1, glucose 25 (Gire & Schoppa, 2008). Slices were aligned on the MEA by locating bands of spontaneous activity (spikes and local field potentials) that indicated the mitral cell and external plexiform layers (Figure 1B; Peace et al., 2018). The MEAs incorporated 120 titanium nitride electrodes (30μm diameter, 30-50 kΩ impedance, 200 μm pitch), embedded within a thin polyimide foil perforated (20-90 μm diameter perforations) to facilitate perfusion and oxygenation of the slice from both sides and to draw the slice down against the MEA with gentle vacuum, thereby improving the signal-to-noise ratio. Recordings were bandpass filtered (1 Hz – 3.3 kHz) and amplified (1200x) before sampling at 20 kHz with a bit depth of 24 bits per sample. This bit depth enabled spike shapes to be recorded at high resolution even at amplifications low enough to also record LFPs.

### Spike sorting

Recorded data were zero-phase (forward + reverse) bandpass filtered between 300 Hz – 3 kHz prior to offline spike sorting and unit identification using the SpykingCircus software package (Yger et al., 2018). Well-isolated candidate units were analyzed as single-unit data when (1) all spike events had amplitudes of greater than 5 standard deviations above the noise of the voltage signal (i.e., thresholded based on the signal to noise ratio of the individual recording), and (2) a refractory period was clearly visible in the interspike interval distribution. Any spikes not clearly separable by these criteria were treated as multiunit data. These conservative spike-sorting criteria were designed to minimize the likelihood of spikes from separate units being incorrectly combined into single-unit data, so as to improve the reliability of subsequent analyses of spike-mediated information.

### Analysis of stimulus-evoked responses

To visualize the instantaneous firing rates of single units, spike trains were convolved with a negative exponential kernel (20 ms time constant) for each trial, and then averaged across trials. Although considerable variability was observed in the time course of unit responses to stimulation, stimulus-responsive units generally displayed changes in firing rate shortly after stimulus onset (Figure 1). To capture these initial responses, unit firing rates were calculated during the first theta/sniff cycle (≈166 ms) of fictive odor stimulation and compared to spontaneous activity measured over the same period prior to stimulus onset. To identify responsive odor-unit pairs, we performed a one-sample t-test (two-tailed, α=0.05) on the stimulus-evoked change in firing rate. Responsive odor-unit pairs were then classified as excitatory (“E”) or inhibitory (“I”) based on the directionality of the change in firing rate. All other odor-unit pairs were classified as non-responsive (“N”). This method of classifying odor-unit responses yielded qualitatively similar results across different values of α.

### Spike-LFP analyses

Recorded data were low-pass filtered at 250 Hz and resampled at 500 Hz for LFP analysis. The Fitting Oscillations and One-Over-F (FOOOF) algorithm then was applied for parametric model-based decomposition of LFP power spectra to identify electrodes with spectral peaks in the gamma band (Donoghue et al., 2020; Supplementary Figure S1). LFP data from electrodes with detectable gamma peaks then were band-pass filtered from 20-55 Hz (third-order Butterworth filter), and the Hilbert transform was applied to extract phase information. To calculate phase-locking values (PLVs), each spike was represented by a unit vector with a phase derived from the concurrent LFP. Then, for a given time window, spike-LFP relationships can be described by a vector sum of all of the spikes occurring in that window; this vector sum is a complex number that contains both phase and amplitude information. The preferred phase for a neuron is determined by the phase of the vector sum, whereas the PLV, which represents the degree to which a particular neuron is synchronized to a particular phase of the LFP oscillation, is determined by the vector sum amplitude divided by the number of spikes during the trial. However, the PLV metric still is known to be biased with respect to the total number of spikes (Vinck, van Wingerden, Womelsdorf, Fries, & Pennartz, 2010). To correct this bias, we applied Rayleigh’s uniformity transformation to obtain a Z-score (*Z = n* × *PLV^2^*, where *n* is the number of spikes) and its corresponding *p*-value (*p = e^−z^*), so as to assess the probability of observing a given PLV by chance given the spike count (Fisher, 1993). Importantly, the *p*-value resulting from this transformation is not being used as a hypothesis test for statistical significance, but as a corrected, standardized, descriptive metric of phase-locking between a spiking neuron and the underlying LFP. Finally, to preclude the possibility of spikes being insufficiently filtered out of LFP recordings and thereby biasing PLV computation, we only compared spikes with LFP phases derived from different electrodes.

### Unitary event analysis

To assess spike-spike synchronization patterns without reference to the LFP, we performed unitary event analyses using disjunct (“fixed”) time bins (Grun, 2009; Grun, Diesmann, & Aertsen, 2002a; Grün, Diesmann, & Aertsen, 2010). The spiking activity of each recorded neuron was represented as a binary matrix with a 5 ms time discretization (in which a one represents the presence of at least one spike in the window, and a zero the absence of any spike). This time discretization reflects an appropriate phase constraint window on the gamma timescale (Figure 3C) – e.g., 5 ms corresponds to one quarter cycle of a 50 Hz gamma oscillation. For each pair of neurons, time bins containing spikes from both neurons in a given trial were defined as coincident events. The empirical coincidence count, *n*_emp_, then was calculated by summing the number of events across the first second of each stimulus presentation trial, across all ten trials. The probability of observing any value of *n*_emp_ can be calculated analytically based on the mean firing rates of each neuron; however, this approach does not account for nonstationarities in firing rate within or across trials (Grun, 2009; Grun, Diesmann, & Aertsen, 2002b) – both of which were present in our data. To account for these nonstationarities, we generated 10,000 surrogate datasets for each spike train on a trial-by-trial basis by randomly dithering each spike within a window of ±10 ms of its original position. Coincident events then were calculated for each surrogate dataset to produce a distribution of expected coincident counts under the null hypothesis (*n*_exp_). The probability of observing *n*_emp_ coincidences within the distribution of *n*exp then was used to evaluate the spike synchrony between neurons. Specifically, this probability was logarithmically transformed into the surprise measure, 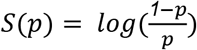, which we used to compare group means and for visualization (Grun, 2009; Palm, Aertsen, & Gerstein, 1988).

## Acknowledgments

We thank the organizers of the Advanced Neural Data Analysis (ANDA) course at the German Neuroinformatics Node (G-Node) for sharing their knowledge of data analysis techniques, which have been implemented in this work.

Supported by NIH/NIDCD grants F31 DC017382 to JCW, and R01 DC014701 and R01 DC014367 to TAC.

---

**Supplementary figure S1.**
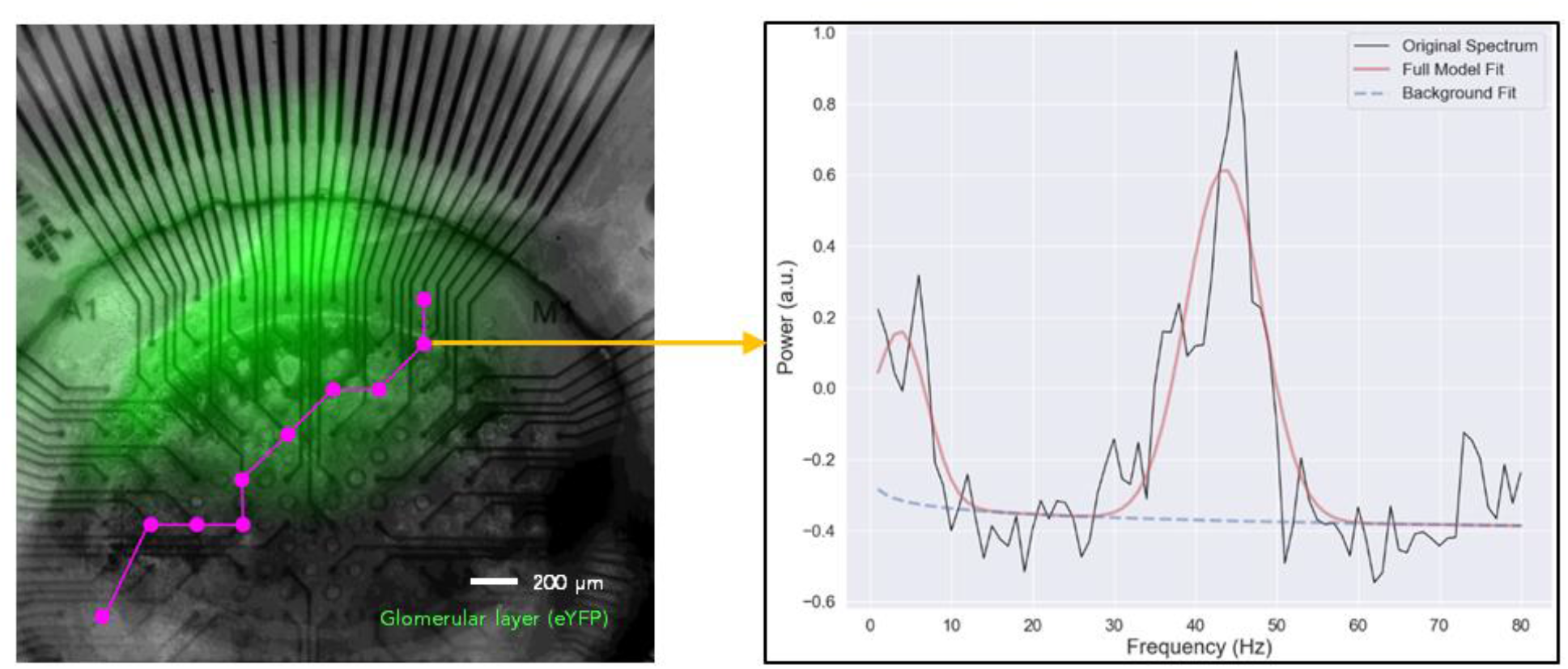
Locations of electrodes across an OB slice exhibiting significant power in the gamma band. *Left:* Horizontal OB slice (50% transparency overlay) laid atop a 120-electrode array expressing EYFP in the glomerular layer (green). Electrodes with detectable spectral peaks in the gamma band are labeled in magenta, forming a band immediately deep to the glomerular layer that presumably corresponds to the EPL/MCL. *Right:* Example of the FOOOF model (Donoghue et al., 2020) used to detect oscillations on each electrode. Presumptive oscillations are identified as periodic components (i.e., spectral peaks) rising above the aperiodic component of the signal (reflecting its 1/f characteristics). Here, a ~42 Hz gamma oscillation is identified. A second, smaller peak at ~6 Hz arises from the 6 Hz sinusoidal modulation of fictive odor input.

**Supplementary figure S2.**
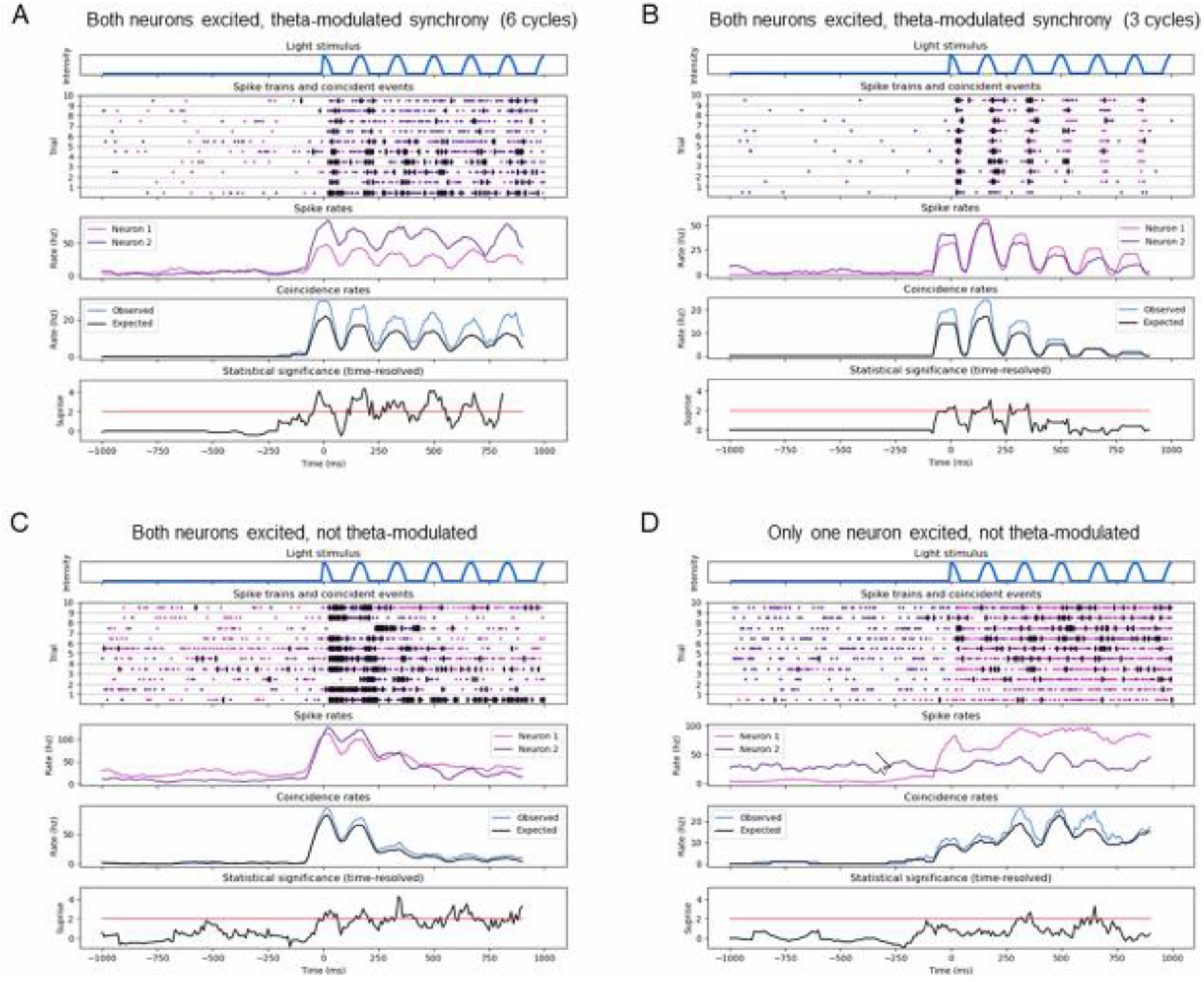
Examples of pairwise synchrony shown with time-resolved UE analysis. For a description of the different plots within each panel, see legend for Figure 4A. **(A)** A pair of neurons that are both excited by the fictive odor, resulting in theta-modulated synchrony for a full second. **(B)** Similar to A, except theta-modulated synchrony only lasts a couple of cycles before returning to baseline levels. **(C)** A pair of neurons that are both activated by a fictive odor input resulting in synchronization. As spike rates return to baseline levels, the synchrony persists, but becomes decoupled from the theta rhythm. **(D)** Only one neuron in the pair is excited by the fictive odor, resulting in weaker synchrony that is not theta-modulated.

